# A major myna problem; invasive predator removal benefits female survival and population growth of a translocated island endemic

**DOI:** 10.1101/2023.04.22.537650

**Authors:** Thomas J. Brown, Max Hellicar, Wilna Accouche, Jildou van der Woude, Hannah Dugdale, Jan Komdeur, David S. Richardson

## Abstract

Invasive predators are a major driver of extinctions and continue to threaten native populations worldwide. Island eradications of (mostly mammalian) invasive predators have facilitated the (re)establishment of numerous island-endemic populations. Other invasive taxa, such as some predatory birds, could pose a more persistent threat due to their ability to fly and actively re-invade even remote and isolated islands. However, the impact of invasive predatory birds has been largely overlooked. We report on a novel sex-specific impact of an invasive-nest predator, the common myna (*Acridotheres tristis*), on a reintroduced population of Seychelles warblers (*Acrocephalus sechellensis*); translocated from Cousin Island to Denis Island in 2004. Regular post-translocation monitoring revealed that female mortality was 20% higher than males, leading to a 60–70% male-biased population sex-ratio between 2005–2015. This was attributed to mynas inflicting severe injuries to incubating female warblers while attempting to prey upon eggs in their nests. These effects likely contributed to the slower-than-expected population growth observed (relative to previous translocations of Seychelles warblers to other islands) over the same period. An eradication programme beginning in 2011 removed all mynas from Denis by 2015. Subsequently, we observed a balancing of sex-specific survival and the population sex-ratio of Seychelles warblers and, consequently, accelerated population growth. This study demonstrates the importance of assessing the threat posed by all invasive taxa (not just mammals) to island conservation. Furthermore, we show how extended monitoring is needed to identify problems, and develop solutions, post-translocation.

## Introduction

The introduction of invasive predators is responsible for almost 60% of contemporary vertebrate extinctions, arguably making them the most damaging animal group for global biodiversity (Doherty *et al*., 2016). Island endemics are particularly susceptible to invasive predators as ecological and evolutionary isolation leaves them ill-equipped to recognize and respond to novel predation, often leading to population declines and local extinctions (Sih *et al*., 2010; Simberloff *et al*., 2013; Carthey and Banks, 2014). For their small surface area, oceanic islands harbour a disproportionate amount of the Earth’s biodiversity, yet have also experienced the fastest rates of species loss (Simberloff, 2000; Fernández-Palacios *et al*., 2021). Understanding and mitigating the impacts of invasive predators on island species is, therefore, crucial for global biodiversity conservation.

Invasive predators are established on the majority of oceanic islands and continue to be one of the most intensive threats to endangered island species (Dueñas *et al*., 2021). Controlling invasive predators can facilitate the re-establishment and growth of native island populations (Jones *et al*., 2016; Prior *et al*., 2018), with perhaps the best examples being the recovery of island bird species following the removal and/or suppression of invasive mammalian predators (e.g. Donlan *et al*., 2007; Lavers, Wilcox and Donlan, 2010; Blanvillain *et al*., 2020). However, since invasive species research tends to focus on species considered to have the most severe impact (i.e. predatory mammals), the impact of other invasive groups can be overlooked. For example, predation of island endemic birds by invasive birds is widespread and, in some extreme cases, have caused extirpations/extinctions (see Evans, 2021). Furthermore, the ability of many invasive birds to actively disperse long-distances means they can (re)colonise islands without anthropogenic assistance (in contrast to most invasive mammals) and thus pose a unique challenge for island species conservation and eradication programs (Feare, 2010; Trakhtenbrot *et al*., 2005). Nonetheless, the impact of invasive birds, and the benefits of their removal, on native species (or: native island species) remain poorly understood (Evans, 2021).

In terms of the victims of invasive predators, eradicating the predator may not always be sufficient in itself. Extirpated island species with limited dispersal capabilities may be unable to naturally recolonize areas of invasive predator exclusion, requiring translocation (the intentional release of a species) to (re)establish populations (Griffith *et al*., 1989; Towns *et al*., 2016). A key determinant of translocation success is ecological suitability at the release site, including the absence of invasive predators (Magdalena Wolf, Garland and Griffith, 1998; Morris *et al*., 2021). For example, translocated population growth can be limited by re-invasions of the release site by invasive predators (e.g. Norbury *et al*., 2014), dispersal of the translocated population beyond areas of invasive predator control (Richardson, Doerr and Parker, 2015; Irwin *et al*., 2021), and contact with new and/or unexpected invasive predator species (e.g. Emery *et al*., 2021). Hence, invasive predators can be responsible for both the need for, and failure of, translocations (Short *et al*., 1992; Bubac *et al*., 2019).

One rarely studied, but potentially important, impact of invasive predators on threatened island populations is an imbalance in the operational sex-ratio; the number of males to females available for mating at any one time. Deviations from a 1:1 operational sex-ratio limits the availability of mates for the more abundant sex – especially in monogamous breeding systems – leading to reduced reproductive output and increased competition among individuals (Nunney, 1993; Le Galliard *et al*., 2005; Heinsohn *et al*., 2019). In small populations, these effects can reduce population growth rate and genetic diversity and, ultimately, increase extinction-risk (Stephens and Sutherland, 1999; Engen, Lande and Sæther, 2003; Bessa-Gomes, Legendre and Clobert, 2004; Jamieson, 2011). Therefore, sex-ratios are an important consideration when conserving and reintroducing threatened species (Komers and Curman, 2000; Wedekind, 2002).

For socially monogamous species, a 1:1 operational sex ratio maximizes the effective population size and growth rate. However, sex-ratios in some taxa, such as birds, tend to be male-skewed, particularly in threatened species and in small and/or declining populations (Donald, 2007; Székely *et al*., 2014; Morrison *et al*., 2016). This pattern has been attributed to females having higher rates of mortality and/or dispersal, with the resulting male-skew restricting population productivity (Steifetten and Dale, 2006; Grüebler *et al*., 2008). In some species, for which females provide the majority of parental care to offspring, a major driver of female mortality is predation during nesting (e.g. Rymešová, Šmilauer and Šálek, 2012; Ramula *et al*., 2018), meaning that increases in predator species abundance, such as the introduction of invasive predators, can increase female (but not male) mortality and cause a male-skewed sex-ratio (e.g. Lehikoinen *et al*., 2008; Stojanovic *et al*., 2014). Further, authors of these studies suggest that reducing female-specific mortality (i.e. removing predators) would disproportionately benefit the recovery of those populations; however, there have been few (if any) empirical tests of this.

Here, we report on the impact of an invasive nest-predator, the common myna (*Acridotheres tristis*, hereafter “myna”), on the establishment of a translocated island population of Seychelles warblers (*Acrocephalus sechellensis*). In 2004, 58 adult Seychelles warblers (of approximately equal sex-ratio) were translocated to Denis Island following the successful elimination of mammalian predators (Richardson, Bristol and Shah, 2006). While individuals started breeding immediately after the transfer, the initial population growth rate was slower compared to previous translocations of the species to other islands (Brouwer *et al*., 2009). Additionally, post-translocation monitoring revealed that females were gaining injuries and had higher mortality rates (compared to males) which, subsequently, resulted in a male-biased adult sex-ratio (unpublished reports: Brouwer *et al*., 2007; van der Woude and Wolfs, 2009; van der Woude and Ploegaert, 2010; van de Woude and Ploegaert, 2011; van der Woude, Apperloo and van Marrewijk, 2013). These observations were attributed to the presence of myna, which were seen attacking incubating female warblers while attempting to prey upon eggs or chicks in the nest. Incubating females do not evade myna attacks despite the risk of injury, as sitting tight on the nest is an evolved response to deter smaller native nest-predators (Veen *et al*., 2000). We hypothesised that the impact of myna predation on reproductive success and sex-ratio were contributing to the slower than expected population growth. An eradication effort from 2011 to 2015 successfully removed myna from the island (Feare *et al*., 2017); however, the effects of this on Seychelles warbler demography had not been assessed. The addition of further data collected in 2022 enabled us to compare rates of sex-specific survival, sex-ratio and population growth before and after the removal of mynas, and to that of myna-free (i.e. control) Islands. This quasi-experimental study provides a rare opportunity to explore the impact of sex-ratio bias, mediated by invasive predation, on island species conservation.

## Methods

### Study system

The Seychelles warbler is a small insectivorous passerine endemic to the Seychelles. Thought to be common and widespread across the Seychelles archipelago historically (Spurgin *et al*., 2014), by the mid-20^th^ century the species nearly became extinct as a result of habitat loss and the introduction of invasive predators. By 1960, only ca 29 individuals remained, occupying a single hectare of mangrove forest on Cousin Island (4.3315° S, 55.6620° E, 29 ha; Loustau-Lalanne 1968). However, with the restoration of native habitat, the Cousin population rapidly increased and stabilized at around 320 individuals from 1980 onwards (Komdeur and Pels, 2005; Brown *et al*., 2021). Seychelles warbler populations are structured into clearly defined territories that are defended year round by a pair-bonded breeding pair, but may also include 1–5 sexually mature subordinates (Richardson, Burke and Komdeur, 2002) which sometimes engage in helping behaviour and cobreeding (Richardson, Komdeur and Burke, 2003; Hammers *et al*., 2019).

Seychelles warblers are reluctant to fly over open water and thus fail to disperse to establish populations on other suitable islands (Komdeur *et al*., 2004). Consequently, a series of translocations were undertaken as part of the species action plan (Richardson, 2001) to establish insurance populations and increase the global population size; which exceeds 3000 across five islands (Wright, Shah and Richardson, 2014). In brief, 29 birds were translocated from Cousin to Aride (4.2129° S, 55.6648° E, 0.68 km²) and Cousine (4.3507° S, 55.6475° E, 0.25 km²) in 1988 and 1990, respectively (Komdeur, 1994). A further 58 birds were translocated to Denis (3.8053° S, 55.6676° E, 1.42 km²) in 2004 and 59 birds to Fregate (4.5837° S, 55.9386° E, 2.19 km²) in 2011, from Cousin (Wright, Shah and Richardson, 2014; Richardson, Bristol and Shah, 2006). These islands were selected for translocations as they contained suitable native-forest habitat and lacked mammalian invasive predators. For all translocations, the founding population had an approximately equal sex-ratio and was released at sites with good quality habitat (i.e. high insect abundance; Richardson, Bristol and Shah, 2006; Wright, Shah and Richardson, 2014; Komdeur *et al*., 1995a)

Common myna have been introduced beyond their native South Asian range to many parts of the world and are considered one of the most damaging invasive species in the world (Lowe *et al*., 2001; Cohen *et al*., 2019). In the Seychelles, mynas predate eggs and interfere with the breeding of endemic bird species, such as Seychelles magpie robins (*Copsychus sechellarum*; Komdeur, 1996). Mynas invaded Denis from nearby islands and, at the time of the Seychelles warbler translocation, a large population of mynas (ca. 1000) had established on Denis (Feare *et al*., 2017). In subsequent years, head injuries (bald patches, scars, and eye damage) were observed on ca. 8% of caught Seychelles warblers and several individuals of other endemic passerine species (van der Woude and Wolfs, 2009), which had not been seen to the same severity or frequency in other Seychelles warbler populations on myna-free islands (i.e. Cousin, Aride, Cousine and Fregate). Video observations of myna birds attacking ‘dummy’ nesting warblers (a taxidermy reed warbler (*Acrocephalus scirpaceus*) sitting on an artificial nest) then confirmed that mynas can inflict such injuries (Woude and Neddermeijer, 2010). This motivated a three-phase eradication, which saw 1186 mynas removed between 2011 and 2015 (see Feare *et al*., 2017 for details). By 2012, ca. 90% of the myna population was removed, with the remaining population removed in 2015.

### Data collection

Post-translocation monitoring of Seychelles warblers on Denis were conducted each year from 2005 to 2015 (with the exception of 2008, 2012 and 2014) and then most recently in April 2022. The duration of monitoring events (hereafter field seasons) ranges from three weeks to two months and involves two or three observers searching the entire island for Seychelles warblers. Individuals are located visually by listening for their songs, calls, or the snapping of their bills while foraging. Individuals are also attracted and located by whistling, ‘pishing’ and using playback of song along paths. Most sighted individuals were followed for ca. 15 minutes to determine territory boundaries and the presence of any associated individuals (e.g., social partner, additional birds and fledglings within the territory; Richardson *et al*., 2007). The movements of individuals and locations of interactions with neighbouring individuals (e.g. fights, alarm calling) were used to infer territory boundaries. Individuals ringed during or before a given field season were visually identified from their unique combination of three colour bands and a British Trust for Ornithology (BTO) metal ring. The probability of re-sighting ringed Seychelles warblers between field seasons on Denis is high (2005 – 2006 = 95%, 2011 – 2013 = 91% and 2013 – 2015 = 82%) and, importantly, does not vary between sexes (Brouwer *et al*., 2007, 2009; van der Woude *et al*., 2013). Furthermore, migration between islands by Seychelles warblers is virtually non-existent (Komdeur *et al*., 2004). Therefore, ringed individuals that are not seen in a given field season can be confidently assumed dead.

During each field season, catching attempts are made throughout the island, particularly in territories/areas with a high proportion of unringed individuals. Individuals are captured using mist nets and conspecific playback (see Kingma *et al*., 2016 for details). Newly caught individuals are given a unique combination of colour bands and a metal ring with unique number (to facilitate future identification), blood sampled and classed as juveniles (<8 months old) or adults based on eye colour, which transitions from grey to red-brown (Komdeur, 1992). The sex of individuals is confirmed molecularly using a PCR-based method (outlined by Griffiths *et al*., 1998) using 25ul of blood drawn with a microcapillary tube from the brachial vein. This procedure is a routine, nonlethal way to sample blood from passerine birds and has been shown to have no measurable impact on condition or survival (Sheldon *et al*., 2008). Furthermore, the monitoring and catching methods outlined are routinely and consistently performed by members of the Seychelles warbler project across all five island populations (Davies *et al*., 2021; Brown *et al*., 2022).

### Population estimates

At the end of each field season, all observations made during visual monitoring and catching are collated and plotted on a map to determine territory boundaries and occupants, i.e., all individuals sighted in territories. From 2004 to 2015, it was possible to perform full censuses due to the very limited population size and high proportion of ringed individuals on Denis. Hence, for this period the sum number of observed individuals, represents the estimated population size for a given field season. In 2013 and 2015, these estimates were corrected following resighting probabilities, which assume 9% and 18%, respectively, of the true population was unobserved. In 2022, we were unable to observe and/or capture the majority of individuals present due to a substantial increase in population size and long length of time since the last period of ringing (2015-2022). Therefore, we estimated population size by extrapolating observed Seychelles warbler densities in comprehensively surveyed areas. We calculated average territory size (hectares) and average group size (number of resident individuals in territories), from a subset (62.5%) of mapped territories for which we had sufficient data to confidently ascertain territory/group size. Using these two values we estimated Seychelles warbler population density (birds/hectare).

Population size was estimated by multiplying estimated density by the total land area of Denis Island (132 ha), excluding unusable areas; beaches, grassland, buildings, and intensified agricultural land (9.65 ha). This population estimate (hereafter ‘maximum estimate’) assumes that density is constant across the island. While our survey confirmed the presence of territories across the whole island (except unusable areas), the frequency of observed territories was lower (*ca* 50%) in coconut palm forest and plantation areas compared to native forest (Fig. S1), which is consistent with prior knowledge that these are less suitable habitats (Johnson *et al*., 2017). Therefore, we produced two more population estimates using the following scenarios: 1) zero occupation of the palm forest/plantation area (26 ha) by Seychelles warblers (hereafter “ultraconservative” estimate), and 2) the density of Seychelles warblers in the palm forest/plantation area is 50% of the estimated density (hereafter “moderately-conservative” estimate).

The population size of myna on Denis was estimated each field season from the beginning to the end of the eradication (2010 – 2015) using transect point counts. The Island was divided into six areas based on habitat type, with five fixed 200m transects in each area. Counts were conducted at five points along 200m transects at 50m intervals. During counts, a single observer counted all mynas sighted within a 25m radius within a two-minute duration. Counts were performed in the morning, with each transect (within the same area) being counted twice in a random order. For each field season, all counts were completed within 4 – 8 days in similar weather conditions. Population size was estimated as the sum of mynas counted per transect (averaged across replicated counts) multiplied by the proportion of the islands area which was surveyed (total island area (132ha)/surveyed area (5.89ha)).

### Statistical analyses

All statistical analyses were performed in RStudio (version 1.2.5033 and R version 4.0.3, Rstudio Team, 2020). Survival, defined as whether or not an individual present in the previous field season was observed (i.e. alive) in the current field season, was fitted as binomial responses (yes vs. no) with a log link function in a Generalized Linear Mixed Model (GLMM) in lme4 (v1.1-25; Bates *et al*., 2015). To test where survival rates differed between males and females, sex was entered as a fixed effect. The time interval between field seasons (hereafter ‘interval’), was not consistent; while the majority were annual, some were biennial (2007–2009, 2011–2013, 2013–2015) and the two most recent field seasons were seven years apart (2015–2022). Because the proportion of surviving individuals decreases with time between field seasons, interval was entered as a controlling fixed effect. Individuals that were present across field seasons had multiple measures of survival; thus, individual identity was included as a random intercept. Similarly, field season (n = 9) was also included as a random intercept to control for non-independence of survival measures (Brouwer *et al*., 2006). In addition to testing whether there is an overall sex-specific difference in survival rates, we were interested in whether sex-specific survival differed across field seasons, with the prediction that female survival will increase and be equal to male survival in the period after the myna eradication compared to before. For this, we ran multiple univariate binomial General Linear Models (GLMs), with survival as a function of sex, for each field season. For field seasons with intervals greater than one year, we report annual survival (i.e., raw survival probabilities per sex adjusted for time (years) between field seasons) for easier interpretation.

We were interested in whether the male bias in the adult population had decreased since the myna eradication. The population sex-ratio for each field season is estimated by the proportion of the caught adult population (for which sex is known) that are males. Although many more individuals were caught and ringed in 2022 (n = 269) this accounted for only ca 30% (269/875) of the estimated population whereas for the 2004–2015 period, 60– 90% of the population were caught via ‘targeted’ catching i.e. (re)visiting territories to catch known unringed individuals. We expect the 2022 catches to be male-biased as males, being more territorial, are more frequently caught using catching methods which involve playing conspecific song to simulate an intruder on the focal birds’ territory. Therefore, rather than comparing the 2022 sex-ratio to that in previous years, we compared contemporary catch rates (male versus female) across populations; Denis 2022, Frégate 2022 and Cousine 2019. If the Denis population sex-ratio remains male-biased, we would expect the probability of catching males to be higher than on Frégate and Cousine; on which an equal sex-ratio is assumed. Frégate and Cousine were chosen for comparison as these field seasons similarly involved catching a sample of individuals from a largely unringed population. Crucially, neither population has been exposed to large populations of mynas, nor have a known history of male-bias (Komdeur *et al*., 1995b), in contrast to Denis. Using a binomial GLM, the sex of catches (excluding juveniles) was regressed against field season (categorical; Denis 2022, Fregate 2022 and Cousin 2018).

To examine population growth (i.e., population size as a function of time) on Denis we used a segmented regression analysis from the segmented package (v1.6-0; Muggeo, 2022). In contrast to a simple linear regression, segmented regression identifies points in the x-y relationship where the slope changes, termed change points. Therefore, segmented regression analysis is suited for investigating the effects of interventions on time-series data (Wagner *et al*., 2002). We predicted that a change point would occur after the myna eradication, with faster population growth compared to before the myna eradication. The Davies test was used to determine whether regression slopes before and after the change point were significantly different. We compared the variance explained by the segmented model to that of competing regression models assuming linear and exponential growth, respectively, using adjusted *R*^*2*^.

## Results

### Sex-specific survival

Since being translocated to Denis and before eradication of mynas (2004-2015), females in the Seychelles warbler population have had lower survival rates than males (mean annual survival probability; females = 0.70 (n = 399), males = 0.86 (n = 692), GLMM-Sex; β = 0.985 ± SE = 0.142, *z* = 6.956, *P* < 0.001; Fig. 1). This is in contrast to the source population on Cousin Island, for which annual survival is 0.85 for both sexes (Hammers *et al*., 2015; Brown *et al*., 2022). When sex-specific survival was analysed separately by year, female survival was significantly lower in all years except 2005, 2006 and 2010. The magnitude of difference between male and female survival was broadly similar between 2005 and 2015 (Fig. 1). In contrast, survival of females and males from 2015-2022 (denoted as 2022 in Fig. 1) were nearly identical (mean annual survival probability; females: 0.87 (n = 69), males: 0.88 (n = 149), GLM-Sex; β = 0.224 ± SE = 0.444, *z* = 0.505, *P* = 0.614), based on 12% of females and 14% of males alive in 2015 being resighted in 2022, reflecting an increase in female survival.

**Figure 1:**
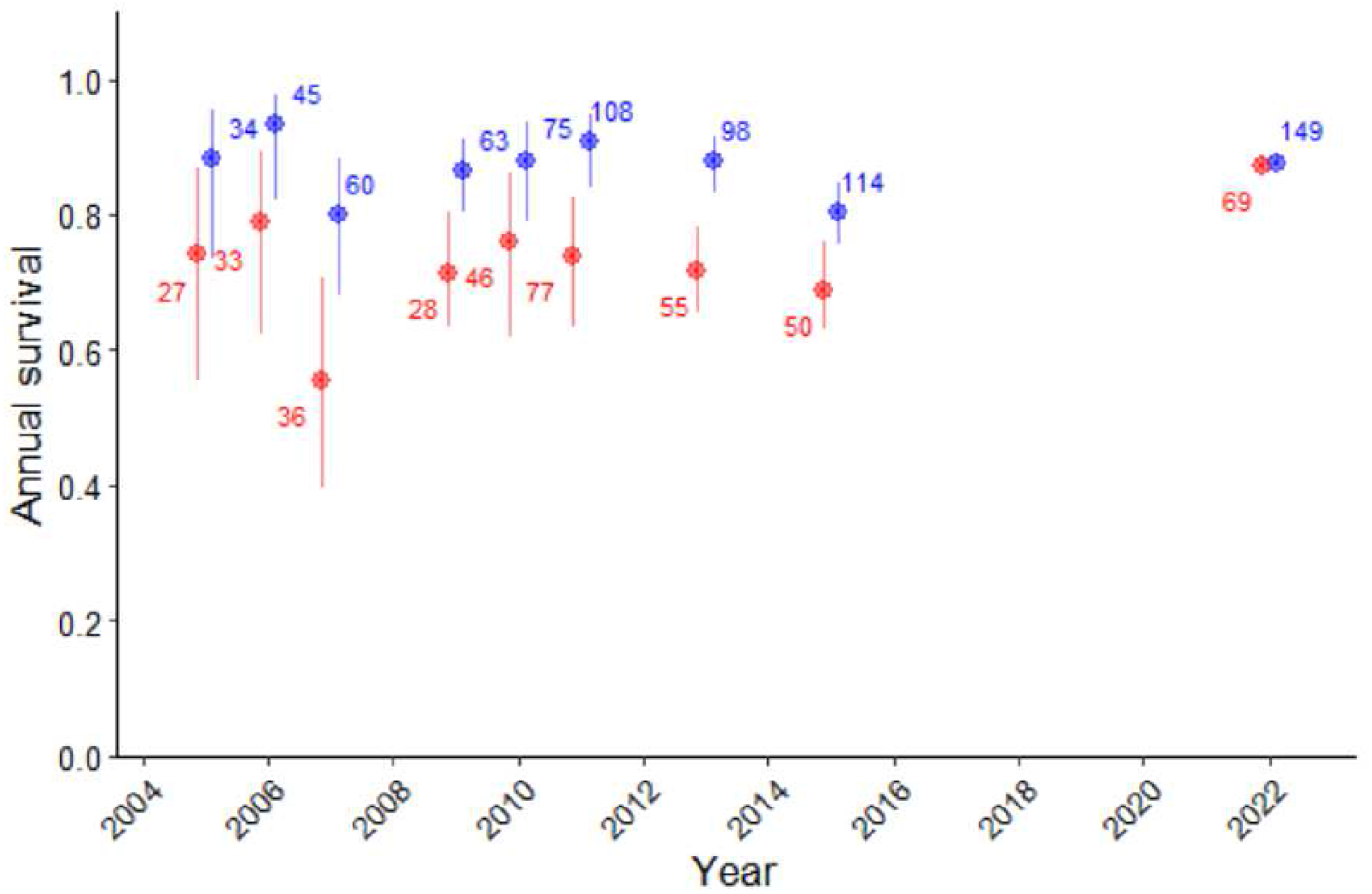
The annual survival probability of Seychelles warblers, based on the proportion of ringed individuals from the previous field season alive in the following field season (x-axis), relative to sex (red = female, blue = male). Points and error bars are mean survival and binomial 95% confidence intervals of raw data, grouped by sex. Numbers refer to sample sizes i.e. ringed individuals present in the previous field season.

### Sex-ratio and catch rates

The Denis adult population sex-ratio has been previously recorded as ca. 60–70% male from 2005– 2015, based on ca. 60–90% of the population having been molecularly sexed in any given field season. Of the 266 individuals newly caught and sexed in 2022, 62% of adults (n = 200) and 48% of juveniles (n = 66) were male, respectively. Of adults caught on Fregate (2022, n = 111) and Cousine (2019, n = 35), 58% and 54% were male, respectively. The probability of an adult catch event being male (versus female) was not significantly different across the three populations (GLM, Predictor = Island (reference = Denis), Fregate; β −0.399 ± SE = 0.239, *z* = −1.668, *P* = 0.095, Cousine; β = −0.318 ± SE = 0.369, *z* = - 0.860, *P* = 0.390), indicating that contemporary adult sex-ratios are similar despite the earlier impact of mynas on Denis.

### Population growth

In 2022, we estimated population density at eight individuals per hectare and a total population size between 771 and 979 individuals (ultra-conservative and maximum estimates, respectively). The moderately-conservative population estimate (assuming 50% density (4 birds/hectare) in the palm forest and plantation areas of the island) of 875 individuals still represents a two-fold (106%) increase from 2015 and an annual growth rate of 15.1%. However, the overall population growth rate (from 2004–2022) is considerably slower than what was initially observed on Aride and Cousine, which appeared to grow very rapidly post-translocation to reach asymptotic densities (i.e., carrying capacity) at ca. 8 and 6 years, respectively (Fig. 2). The initial population growth rate on Denis is more comparable to that observed on Fregate (Fig. 2).

**Figure 2:**
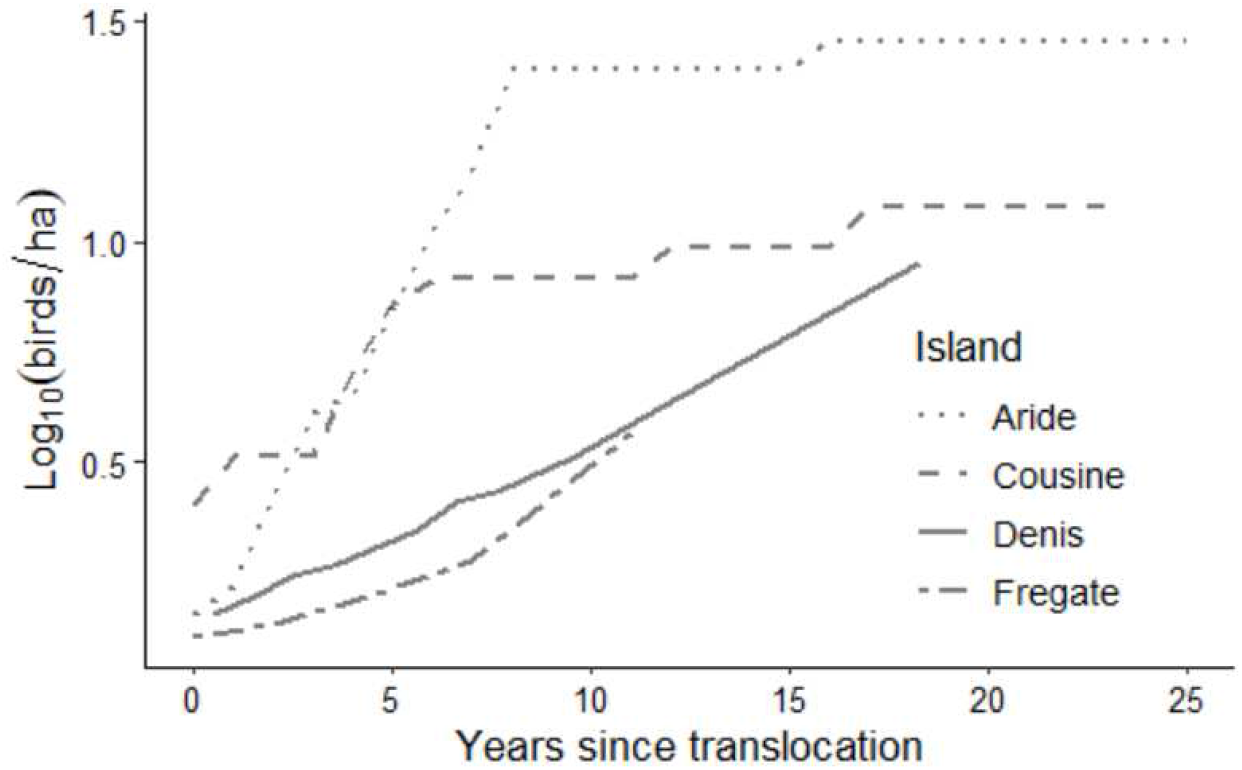
Seychelles warbler population density estimates in the years shortly after translocation to Aride (1988), Cousine (1990), Denis (2004) and Fregate (2011), respectively.

The segmented regression model estimated a change point at 2013, with a significantly faster increase in population size occurring after 2013 compared to before (Davies test; *P* < 0.001; Fig. 3). This is 1.5 years after the first phase of the myna eradication, in which ca. 90% of the myna population was removed (Fig. 3). The pre-change point regression slope, describing population growth from 2004– 2013, predicts a population size of 467 by 2022; 400 individuals less than our moderately-conservative estimate (Fig. 3). The segmented model had the highest adjusted *R*^*2*^ (0.99) followed by the exponential and linear regression models (0.98 and 0.91, respectively). Therefore, observed population growth is best described by models depicting an accelerated increase over time.

**Figure 3:**
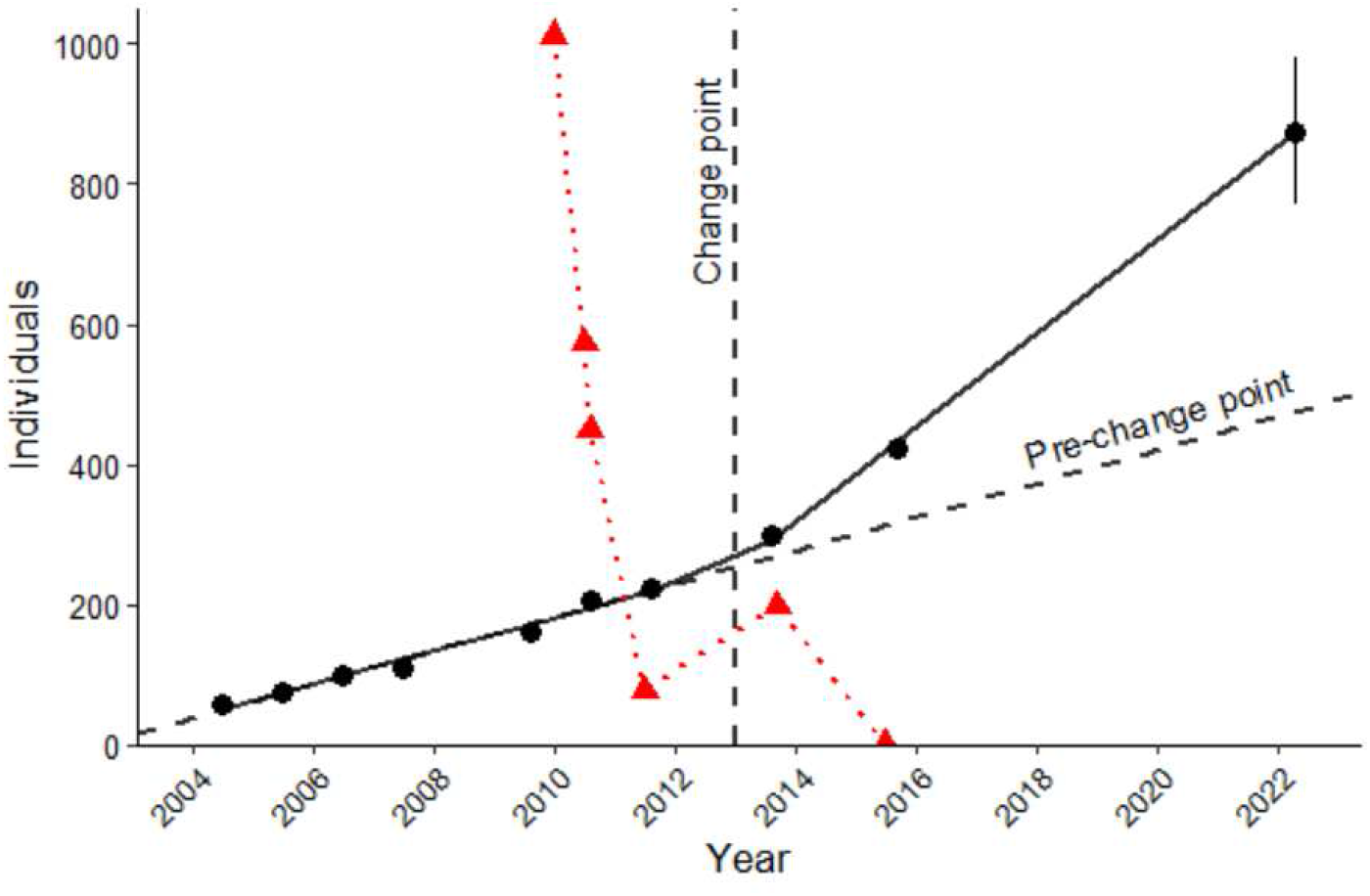
Population size estimates for Seychelles warblers (black) and common myna (red) on Denis Island relative to time (year). The solid fit line for Seychelles warblers is the segmented regression prediction. The dashed lines represent the change point of the segmented relationship (2013) and the continuation of the population growth rate before the change point. The error bar at 2022 depicts the minimum and maximum range of the estimated population size at this point.

## Discussion

We found that the translocated population of Seychelles warblers on Denis Island had higher rates of female mortality compared to male mortality across much of its existence. A notable exception was the period after which myna were removed from the island (2015–2022), when female mortality had decreased to become near identical to male mortality. Of adult individuals newly caught in 2022, 61% were male. This appears to reflect a male-bias in our catching methods rather than a male-biased population, as similar male catch rates are achieved when sampling two other populations. This suggests that, in line with equal male-female mortality, the population sex ratio is approximately equal after the removal of the mynas. The rate of post-translocation population growth on Denis was initially slower than that observed on other (myna-free) islands. However, since the removal of mynas from Denis, population growth has accelerated.

Compared to source and/or resident populations, higher mortality in translocated populations may be expected for a variety of reasons, including new predation pressures (Bradley *et al*., 2012; Parker *et al*., 2013; Norbury *et al*., 2014). However, our study is, to our knowledge, unique in showing that post-translocation mortality and predation-pressure can be sex-dependent. Seychelles warblers (both males and females) on Cousin Island and males on Denis have very similar mortality rates; higher mortality was only observed in Denis females prior to the myna eradication. The most plausible explanation is myna nest-predation, which can inflict life-threatening injuries to parent birds (van der Woude, Apperloo and van Marrewijk, 2013). As for many taxa, Seychelles warbler females spend more time at the nest – only females incubate and have higher nestling provisioning rates – than do males (van Boheemen *et al*., 2019), and thus are more vulnerable to myna attacks. Dispersal by warblers into anthropogenic habitats (i.e. agricultural and recreational; van der Woude and Wolfs, 2009) soon after translocation may have increased exposure to myna-predation, as these habitats typically support higher densities of mynas compared to native forest (Grarock *et al*., 2014; Feare *et al*., 2017). Thus, our study supports the idea that mortality in the more vulnerable sex will increase as small populations disperse into less suitable habitat (Dale, 2001; Richardson, Doerr and Parker, 2015).

Despite mynas being listed as being one of the world’s worst invasive species, empirical evidence of their negative impacts on native species is lacking (Lowe *et al*., 2001). Previous studies have shown that establishment of mynas is associated with a decline in the abundance of native bird species (Grarock *et al*., 2012; Colléony and Shwartz, 2020), with competition (for food and nest cavities) and nest interference/predation being the assumed proximate causes (Blanvillain *et al*., 2003; Grarock *et al*., 2012). While numerous myna-predation events (of native bird species) are documented, these normally concern egg and/or chick predation (Evans, 2021). Our study is the first to demonstrate that nest predation attempts by mynas can impact adult mortality at a population-level and thus limit population productivity more than expected from nest-predation alone. In the context of vulnerable island species conservation, our study warns that the severity of mynas as an invasive species may be greater than previously thought (Evans *et al*., 2018; Evans, 2021).

The tendency for small and isolated populations to have biased sex-ratios is well-documented, particularly a male-bias in bird taxa. While this is often interpreted as a consequence of higher reproductive costs in females, obtaining strong evidence – such as higher mortality in females – can be difficult (Dobson, 1987; Morrison *et al*., 2016). Male-skew in small isolated populations can also be caused by female-biased dispersal, and thus the tendency of females to leave populations, rather than female-biased mortality (Dale 2001). However, this is not case in the Seychelles warbler as dispersal is confined within our study area; thus, it is a reasonable assumption that the male-biased sex-ratio (from 2005–2015) was caused by female-biased mortality. Given that the majority of island reintroductions are of species with limited dispersal capabilities, sex-biased mortality is likely to be the primary driver of sex-ratio bias in translocated populations more generally.

Biased operational sex-ratios, in particular male-bias, have been implicated as a major limiting factor of population growth (Le Galliard *et al*., 2005; Steifetten and Dale, 2006; Eberhart-Phillips *et al*., 2017). Likewise, the male-biased Denis population exhibited slower population growth compared to previous translocations to Aride and Cousin (Johnson *et al*., 2017). However, population growth increased after the myna eradication, which suggests that the proportion of unpaired adult males has since decreased (Steifetten and Dale, 2006; Brooke *et al*., 2012). Likewise, our comparison of contemporary sex-specific catch rates suggest that the Denis population has become less male-biased. We cannot exclude the possibility that other island-specific factors (e.g. area, habitat quality) contributed to differential population growth. These factors may explain why Fregate, being larger both in terms of absolute size and the proportion of unsuitable habitat, also exhibited relatively slow growth in population density (Johnson *et al*., 2017). Nevertheless, our results indicate that myna-related impacts (i.e. nest-failure, female-mortality and male-biased sex ratio) play an important role. Given that the success of translocations is typically quantified by population growth rate (Morris *et al*., 2021), our study reiterates the importance of managing factors that can distort population sex-ratio.

Managing and/or removing invasive predators is an expensive and time consuming enterprise, hence the importance of gathering strong evidence as to the benefits of such efforts (e.g. Jones *et al*., 2016). Eradication programs have successfully removed mynas from three islands (including Denis) in the Seychelles, with the aim of facilitating the recovery and/or (re)establishment of threatened native bird populations (Canning, 2011; Feare *et al*., 2017). However, empirical evidence on the success of this particular action is currently limited, circumstantial, or confounded by other conservation action; for example, when myna removal coincides with the removal of other more damaging invasives (e.g. rats) from the same area (Blanvillain 2020; Tindall 2007). Population growth of Seychelles paradise flycatchers (*Terpsiphone corvina*) and Seychelles magpie robins on Denis (both introduced in 2008 and susceptible to myna attacks) also increased after the myna removal, suggesting a positive effect on the establishment of those populations (Feare, Bristol and Van De Crommenacker, 2022). Crucially, our study of Seychelles warblers goes further in showing that i) causes of low population productivity (i.e. female-mortality and male-biased sex-ratio) have lessened post-removal, and ii) a statistically-supported change-point in the population trajectory, coinciding with the removal, after which growth was significantly faster. Therefore, our study provides the strongest evidence yet that controlling mynas can benefit the re-establishment of native island birds.

## Conclusions

Invasive predators are a major driver of both island species decline and failures to increase the population size/range of island species. Care should be taken to determine the impact of novel invasive predation-pressures – not just those implicated in the initial decline of an island species – on efforts to re-establish island populations. Most post-translocation monitoring is short-term (1 – 4 years) and usually only concerned with translocation success; whether or not the population grows to an adequate size (Bubac *et al*., 2019). However, we show that predation (or attempted predation) can have subtle (but important) effects, such as causing suboptimal population structure and growth rate, which are only detectable with detailed and extended post-translocation monitoring. Therefore, our study demonstrates the importance of such monitoring for informing follow-up action (e.g. invasive predator control) and for promoting the success of future translocations of the same and similar species.

## Acknowledgements

We sincerely thank the Mason family for their hospitality and warm welcome on Denis Island during our visits. Sincere thanks and appreciation go to the Green Islands Foundation, and to Denis Island management, for facilitating all Seychelles warbler surveys, as well as providing transport and accommodation. This study would not have been possible without the contribution of many fieldworkers, laboratory technicians, students, and database managers during the whole study period. The long-term Seychelles warbler study was funded by the Natural Environment Research Council (NERC) grants to D.S.R. and H.L.D. (NER/I/S/2002/00712, NE/I021748/1, NE/F02083X/1, NE/K005502/1), and NWO grants from the Dutch Science Council (NWO) to J.K., with D.S.R. and H.L.D. (NWO-ALW 823.01.014, 854.11.003). J.W. was funded by a NWO-VICI grant (86503003) awarded to J.K.

