## supplements for "A major myna problem; invasive predator removal benefits female survival and population growth of a translocated island endemic"

Supporting information

**Table S1**: General Linear Mixed Model (GLMM, family = binomial) explaining variation in Seychelles warbler survival (yes/no) across field seasons relative to sex and the time interval between field seasons. Model includes survival data for the 2022 field season.

| **Predictor** | **β** | ***SE*** | ***z*** | ***p-value*** |
| --- | --- | --- | --- | --- |
| (Intercept) | 1.346 | 0.225 | 5.977 | <0.001 |
| Sex (reference = female) | 1.024 | 0.152 | 6.746 | <0.001 |
| Field season interval (years) | -0.602 | 0.078 | -7.740 | <0.001 |
| **Random effect** | **1167 observations** | **Variance** |  |  |
| Individual ID | 514 individuals | < 0.001 |  |  |
| Field season ID | 9 field seasons | 0.132 |  |  |

**Table S2**: Univariate General Linear Models (GLMs, family = binomial) testing whether survival differed between males and females for each field season (significant differences are in bold).

| **Field season (year)** | ***N* (individuals)** | **β (Sex, reference = female)** | ***SE*** | ***z*** | ***p-value*** |
| --- | --- | --- | --- | --- | --- |
| 2005 | 61 | 0.965 | 0.690 | 1.399 | 0.162 |
| 2006 | 78 | 1.327 | 0.734 | 1.808 | 0.071 |
| **2007** | **96** | **1.163** | **0.466** | **2.499** | **0.013** |
| **2009** | **91** | **1.283** | **0.476** | **2.697** | **0.007** |
| 2010 | 121 | 0.835 | 0.496 | 1.684 | 0.092 |
| **2011** | **185** | **1.235** | **0.422** | **2.929** | **0.003** |
| **2013** | **153** | **1.382** | **0.359** | **3.846** | **<0.001** |
| **2015** | **164** | **0.917** | **0.349** | **2.630** | **0.009** |
| 2022 | 218 | 0.224 | 0.444 | 0.505 | 0.614 |

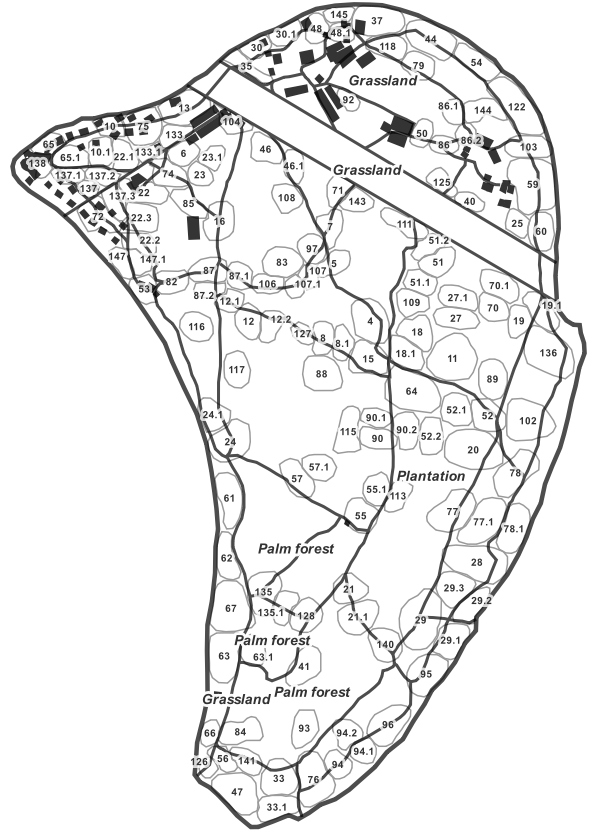

**Figure S1**: 2022 Seychelles warbler territory map of Denis Island.
